# Adding software to package management systems can increase their citation by 280%

**DOI:** 10.1101/2020.11.16.385211

**Authors:** Vahid Jalili, Dave Clements, Björn Grüning, Daniel Blankenberg, Jeremy Goecks

## Abstract

A growing number of biomedical methods and protocols are being disseminated as open-source software packages. When put in concert with other packages, they can execute in-depth and comprehensive computational pipelines. Therefore, their integration with other software packages plays a prominent role in their adoption in addition to their availability. Accordingly, package management systems are developed to standardize the discovery and integration of software packages. Here we study the impact of package management systems on software dissemination and their scholarly recognition. We study the citation pattern of more than 18,000 scholarly papers referenced by more than 23,000 software packages hosted by Bioconda, Bioconductor, BioTools, and ToolShed—the package management systems primarily used by the Bioinformatics community. Our results suggest that there is significant evidence that the scholarly papers’ citation count increases after their respective software was published to package management systems. Additionally, our results show that the impact of different package management systems on the scholarly papers’ recognition is of the same magnitude. These results may motivate scientists to distribute their software via package management systems, facilitating the composition of computational pipelines and helping reduce redundancy in package development.

**Significance Statement:** Software packages are the building blocks of computational pipelines. A myriad of packages are developed; however, the lack of integration and discovery standards hinders their adoption, leaving most scientists’ scholarly contributions unrecognized. Package management systems are developed to facilitate software dissemination and integration. However, developing software to meet their code and packaging standards is an involved process. Therefore, our study results on the significant impact of the package management systems on scholarly paper’s recognition can motivate scientists to invest in disseminating their software via package management systems. Dissemination of more software via package management systems will lead to a more straightforward composition of computational pipelines and less redundancy in software packages.

---

Quantitative evaluation of scientists’ scholarly contributions plays a prominent role in their hiring, promotion, tenure, funding, or institutional rankings [1–3]. Therefore, there has been an increasing interest in the “science of science,” where bibliometric and scientometric analyses are performed to predict outcomes and suggest career trajectories [4,5].

The number of citations a publication receives has been the dominant quantitative recognition of scholarly contribution [4], which is used as a proxy for most impact metrics [6]. Therefore, there has been a proliferation of quantitative evaluation of scholarly literature to identify driving factors in a scholarly paper’s citation count. A myriad of factors have been identified to be involved in recognition of a scholarly paper; some of the widely reported ones are: (i) A phenomena are commonly known as “rich-get-richer” or Matthew effect [7] where highly cited papers are more likely to be cited again [8–12]; (ii) Aging, where the impact and novelty dilute w.r.t. newer works, excluding *sleeping beauties* whose relevance is not recognized for decades [4,13–15]; (iii) Author reputation impact [16,17]; (iv) Novelty, where those scientific discoveries developing new ideas leveraging established elements are generally better received than new or developmental research [4,18,19]; and (v) Team size, where publications of larger teams which generally discuss developmental work are more cited than those of smaller teams with more disruptive studies [20].

A growing number of methods and protocols discussed in scholarly papers are being disseminated as open-source software packages [21]. Here we study the impact of the software packages’ accessibility and reusability on recognizing their respective scholarly papers.

The software packages are used as building blocks in other studies. In addition to the novelty and impact of the operation a package performs, its adoption is a function of various factors, including its discoverability and ease of integration with other packages. However, with the packages being developed by different teams makes them discoverable and installable through various means and not necessarily easily integrated with other packages. Accordingly, several software package management systems (PMS) have been developed to implement a unified interface for searching, installing, and integrating packages, making them more accessible and reusable.

We assess PMS’s role in facilitating software dissemination by comparing the pattern of the citations the software packages receive before and after they were added to a PMS. We study software packages available from Bioconda [22], Bioconductor [23], BioTools, and ToolShed [24]—the PMS’s used by the Bioinformatics community—their characteristics are summarized in Table 1. The study is based on more than 11,000 publications of more than 23,000 software packages.

**Table 1.** A comparison of the characteristics of four software package management systems whose effect in making packages more discoverable is studied here.

| PMS | Target Platform | Contains Executables | Install Packages | Quick Package Documentation | Package Code Review | Continuous integration | How are packages added? |
| --- | --- | --- | --- | --- | --- | --- | --- |
| Bioconda | Conda | ✓ | ✓ | ✗ | ✗ | ✗ | By Developer |
| Bioconductor | R programming language | ✓ | ✓ | ✓ | ✓ | Nightly builds | By Developer |
| BioTools | NA | ✗ | NA | ✗ | NA | ✓ | Automatic |
| ToolShed | Galaxy | ✓ | Galaxy-specific | ✓ | ✗ | ✗ | By Developer |

Our results suggest that the citation count of the publication where the package is discussed increases by up to 280% after the package is added to a PMS—hence underscoring a positive correlation between a package’s availability from PMS and its publication’s citation count. Additionally, our study suggests that the citation count of publications of software packages is boosted twice: once when their manuscript is published, once when the software is added to a PMS, where in some cases the effect of the latter is more significant than the former.

Some factors affecting citation count are inherent (e.g., author reputation and aging), and some can be achieved with minimal effort (e.g., Tweeting [25]); among them, adding a package to a PMS may require a significant amount of effort. For instance, Bioconductor is committed to providing high-quality R packages. Hence the source code of every package is carefully reviewed before the package is added to Bioconductor. Therefore, the quantitative evaluation presented here provides a substantial incentive to devote time to making software packages available from a PMS. Ultimately, this leads to having more packages available from PMS’s, which in addition to attracting more citations, facilitates finding and using packages for specific studies, hence reducing redundancy in the developed packages. In the long-term, it can benefit a community where efforts are devoted to developing new software or improving existing ones instead of developing numerous software with highly overlapping functionalities.

## Results

The present study quantifies the impact of disseminating a software package via PMS’s on their adoption and scholarly paper’s recognition. The impact is quantified by a longitudinal analysis of scholarly paper’s citation count—a proxy metric to quantify adoption and recognition—before and after their respective software package was published to a PMS. The study is focused on software available from PMS’s developed for the Bioinformatics community: Bioconda [22], Bioconductor [23], BioTools, and ToolShed [24]; their characteristics are summarized in Table 1 and detailed in the Discussion section.

The bibliography information of all 23,083 tools available on the PMS’s is extracted and searched on Scopus (see Method section). Among the 18,140 referenced publications, 11,231 are indexed by Scopus (see Figure 2 and Supplementary Figure 1), whose citation per year was retrieved. After normalization (see Method section), these citations are used to study the impact of publishing software to package management systems.

**Figure 1.**
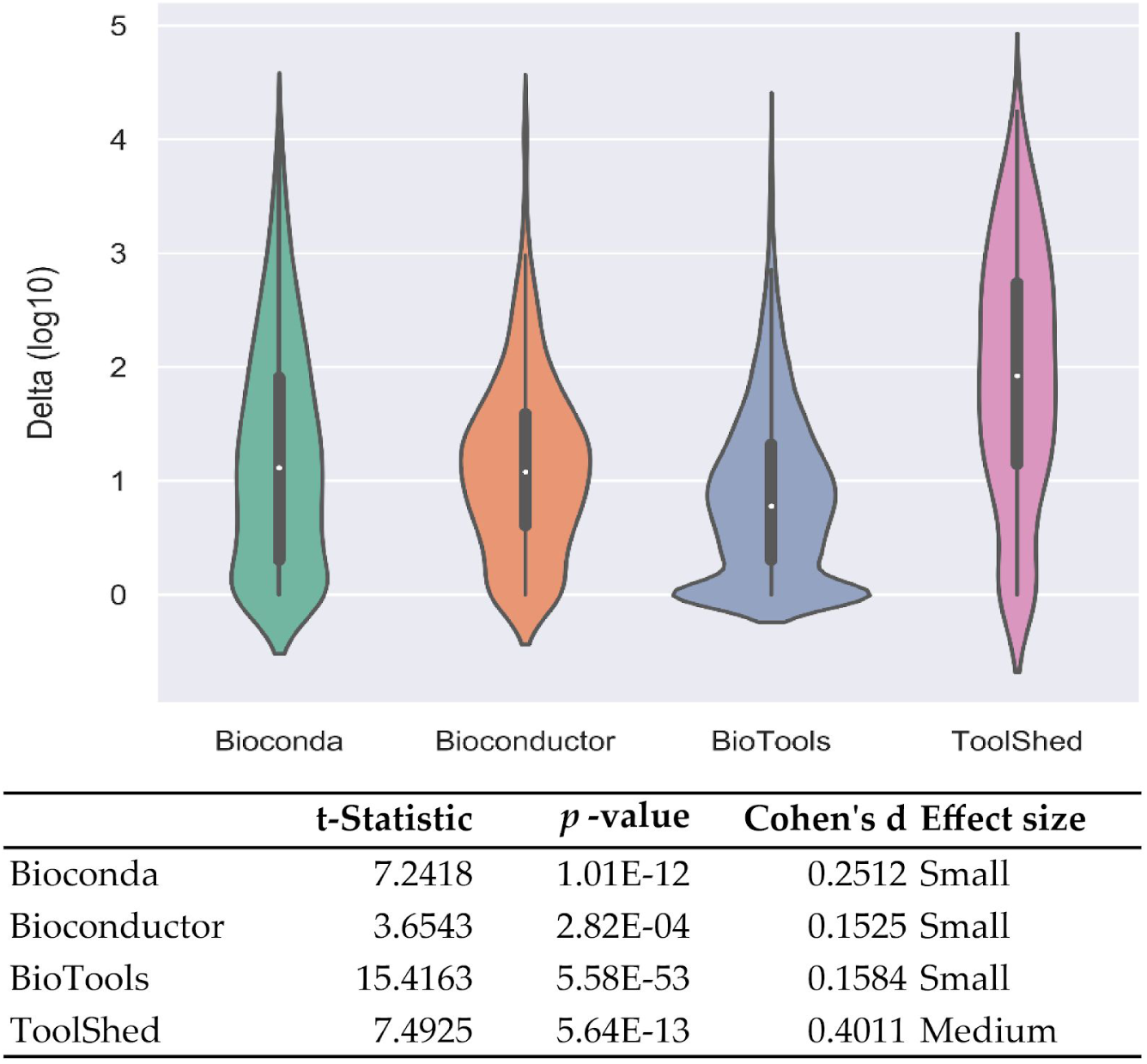
The results of a one-sided one-sample *t*-test for the null hypothesis that the citation count of publications have not changed after their respective software was published to a PMS. In other words, the null hypothesis is that the mean of differences between the sum of the number of citations a publication receives per year before and after its respective software was published to a PMS (*delta*) is zero. The *p*-values suggest that the null hypothesis’s departure is statistically significant for software in all PMS’s. However, Cohen’s *d* suggests that the mean of differences between the two groups of citations in ToolShed is more extensive than zero compared to Bioconda, Bioconductor, and BioTools.

**Figure 2.**
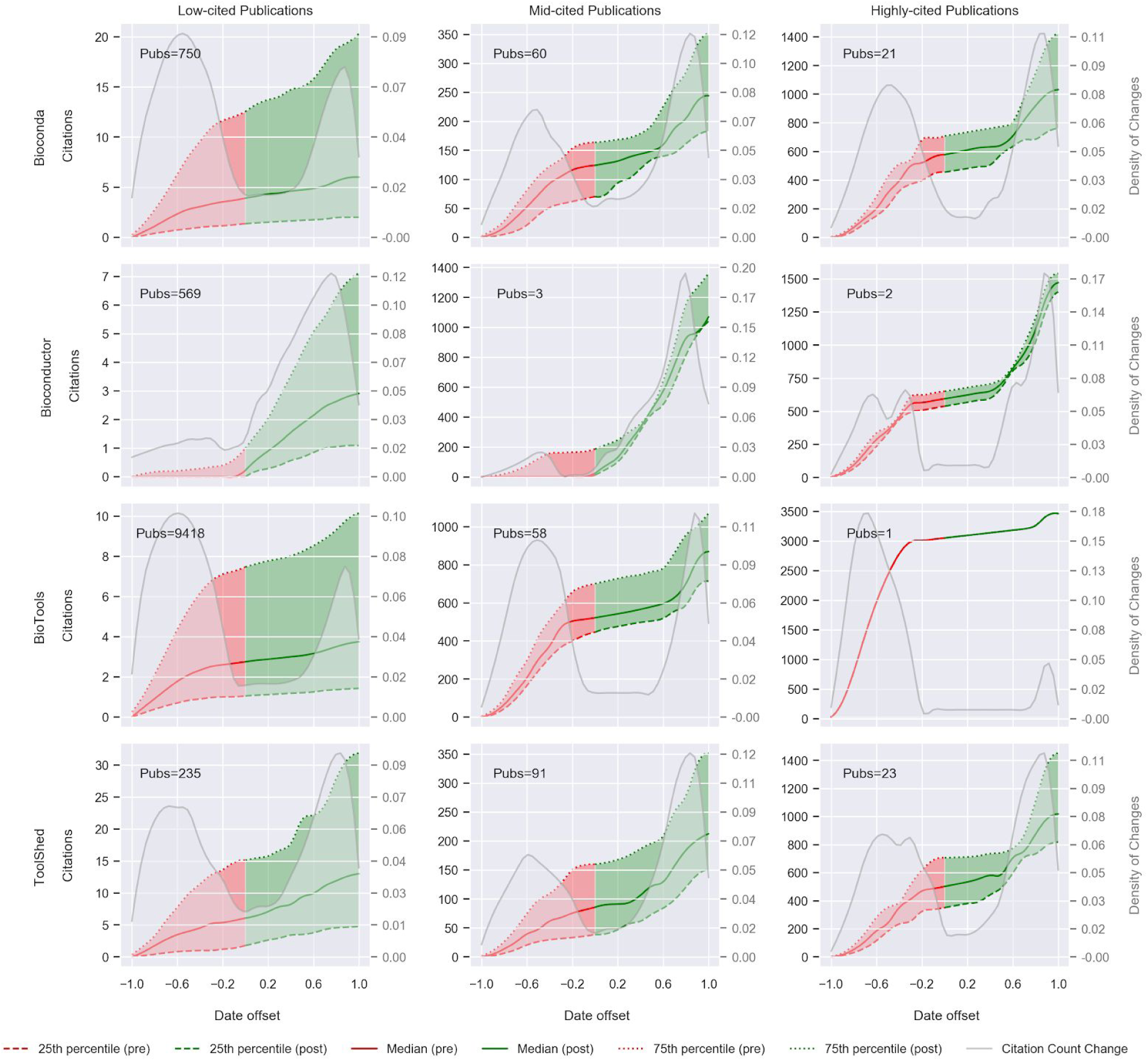
The figure plots the (normalized per year) citation count of publications before (negative date offset), and after (positive date offset), their respective software was added to PMS’s. The publications are grouped in low, mid, and highly-cited clusters (see supplementary Figure 2). The difference between two consecutive citation counts is plotted as *Citation Count Change*, which suggests that their count has boosted twice. Once before, the software was added to PMS’s that can reflect the initial manuscript visibility. Once after, they were added to the PMS’s that reflect the positive influence of being hosted by the PMS’s. The citation increase for packages in Bioconductor does not follow the same pattern as those in other PMS’s, which can be explained using the difference in patterns of when tools are added to repositories w.r.t. their publication date (see Figure 3).

### Citation count increases after the software are published to a PMS

Comparing the number of citations papers received before and after their respective tools were published to PMS’s is an intuitive metric to quantify the impact of disseminating software via the PMS’s on their citation count. For instance, the citation rate of BLAST+ [26] on average was 77.8 per year from 2009 when it was published until 2014 when its respective software was added to ToolShed; the rate then increased to ∼703 citations per year. In addition to ToolShed’s impact, such an increased citation rate can be associated with several driving factors such as “rich-get-richer.” Therefore, to deconvolute the impact of publishing to PMS’s on citation count from other factors, we assert if the difference between the sum of citations a scholarly paper receives per year before and after its respective software was published to a PMS is significantly large to have happened by chance. Accordingly, a one-sided one-sample *t*-test is performed on the difference between before and after citation counts with the null hypothesis that the population’s mean equals zero. The results of the *t*-test are given in Figure 1. The *p*-value and Cohen’s *d* effect size are also reported for each test; the *p*-value quantifies how significantly the null hypothesis is rejected. Cohen’s *d* is a standardized difference that quantifies the magnitude of difference between the means of the two groups of citations.

The tests reject the null hypothesis, suggesting that scholarly papers’ citation count increases after their respective software was published to PMS’s. However, a small effect size of Cohen’s *d* for Biconda, Bioconductor, and BioTools suggests that their means are only slightly above zero. In comparison, it is further above zero in ToolShed. A significant *p*-value with small Cohen’s *d* effect size can reflect many publications with zero citations that inflate the significance. Therefore, in the following section, publications are grouped in different clusters based on their citation count and separately studied longitudinally.

### Clusters of publications

The impact of disseminating software via PMS’s on their adoption may vary depending on the software’s popularity before being added to PMS’s. In other words, publishing software to a PMS may have less effect on highly popular software visibility than the least popular ones. Therefore, we group publications in *high, mid*, and *low*-cited clusters based on their vectors of citations per year using agglomerative hierarchical clustering, which groups most publications in the low-cited clusters (see Table 2). The number of clusters is determined using the *elbow* method [27] that yields a large mean *Silhouette coefficient* for all the clusters (see Method section and supplementary Figure 2). We perform a one-sided one-sample *t*-test on the difference between before and after citation counts of publication in each cluster. The null hypothesis is that the mean of the population equals zero (see Table 2). The tests reject the null hypothesis with a significant *p*-value for low-cited publications with a medium Cohen’s *d* effect size, indicating that the citation count of low-cited publications slightly increases after their respective software was published to PMS’s. The *p*-value of the high and mid-cited publications is also significant (excluding Bioconductor, whose permissive *p*-value is a result of very few publications it has in this group); however, their Cohen’s *d* effect size is “very large,” that indicates relatively popular tools receive a significant citation boost after their respective software was published to PMS’s.

**Table 2.** Results of a one-sided one-sample *t*-test for the null hypothesis that the citation count of publications do not change after their respective software was published to PMS’s. The results indicate that all publications’ citation count increased after their respective software was published to PMS’s. In contrast, the boost in citation count is significantly more considerable for relatively popular publications. The test is not performed for the highly-cited publications of BioTools, because this group contains only one publication.

|  | Cluster | #Publications | t-Statistic | *p*-value | Cohen's *d* | Effect size |
| --- | --- | --- | --- | --- | --- | --- |
| Bioconda | Low | 750 | 12.827 | 3.46E-34 | 0.468 | Medium |
|  | Mid | 60 | 8.077 | 4.05E-11 | 1.043 | Very large |
|  | High | 21 | 6.507 | 2.42E-06 | 1.420 | Very large |
| Bioconductor | Low | 569 | 7.159 | 2.53E-12 | 0.300 | Small |
|  | Mid | 3 | 4.320 | 4.96E-02 | 2.494 | Very large |
|  | High | 2 | 8.514 | 7.44E-02 | 6.020 | Very large |
| BioTools | Low | 9,418 | 24.333 | 6.80E-127 | 0.251 | Small |
|  | Mid | 58 | 11.060 | 8.22E-16 | 1.452 | Very large |
|  | High | 1 | - | - | - | - |
| ToolShed | Low | 235 | 9.589 | 1.40E-18 | 0.626 | Medium |
|  | Mid | 91 | 9.475 | 3.54E-15 | 0.993 | Very large |
|  | High | 23 | 5.659 | 1.09E-05 | 1.180 | Very large |

### Citation counts are boosted twice

A manuscript is publicized at its publication through various means that bring it to the scientific community’s attention. Therefore, manuscripts receive an initial boost in citation count, which flattens as they age [4,14]. Several factors can delay the flattening, such as early citers [17]; however, excluding *sleeping beauties* [14], no factors have been reported to stimulate a citation boost after their ascent has been flattened.

The citation count of 11,231 scholarly papers whose respective software was disseminated through Bioconda, Bioconductor, BioTools, ToolShed is plotted in Figure 2. The plots suggest a citation count of manuscripts is boosted after their publication, which is in accordance with the literature. Additionally, the plots show a second boost in citation count after the software was published to the PMS’s. The slope between two consecutive citation counts shows that the two boosts are of the same scale, underscoring the positive impact of software’ accessibility and discoverability on their adoption.

The manuscripts whose respective software was published to Bioconductor do not follow the same pattern as those hosted by Bioconda, BioTools, and ToolShed. The citation count of these publications is primarily boosted once after their respective software was published to Bioconductor. This pattern reflects the norms the Bioconductor community has adopted where, in most cases, the manuscript and its respective software are published one year apart at the latest (see Figure 3).

**Figure 3.**
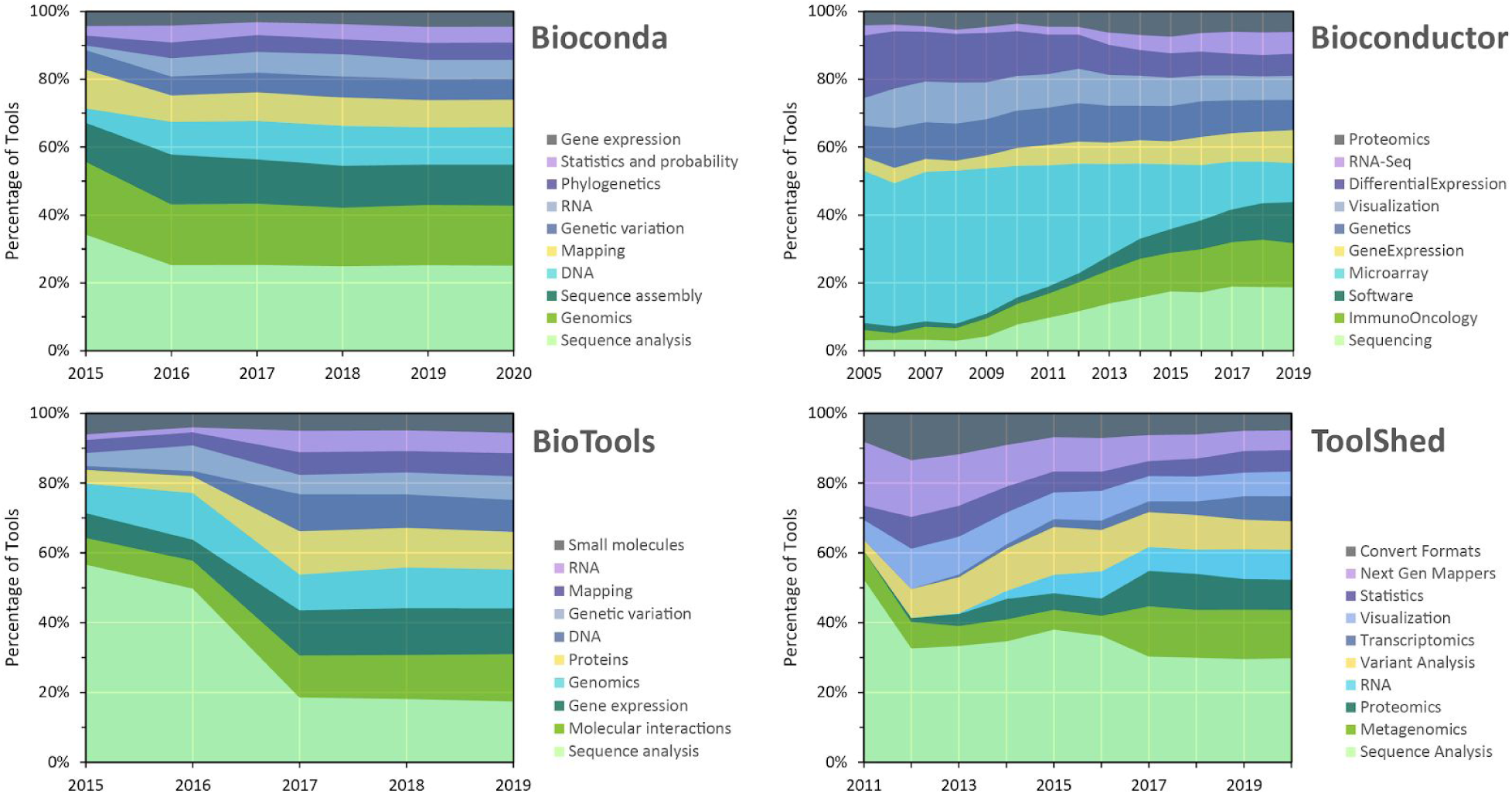
The figure plots the number of tools in the ten most popular categories in 2020 throughout the PMS’s lifetime.

### Any difference between PMS’s?

Bioconda [22], Bioconductor [23], BioTools, and ToolShed [24] are the PMS’s whose hosted software packages are studied here. The PMS’s differ in their characteristics, targeted community, and their impact on adopting the packages they host. The differences are characterized in the following.

#### Differences in the packages they host

The characteristics of these management systems are summarized in Table 1; in general, the packages they host cover a broad spectrum of biomedical data analysis, including genomics, proteomics, and immuno-oncology (see Figure 3). However, they differ in their targeted community, application, and the number of packages they host, and the degree of overlap among them.

#### Differences in their community norms

Generally, authors upload software to PMS’s after their scholarly paper is published, except for the Bioconductor community, where most packages are uploaded at the latest one year apart from their scholarly paper’s publication (see Figure 4).

**Figure 4.**
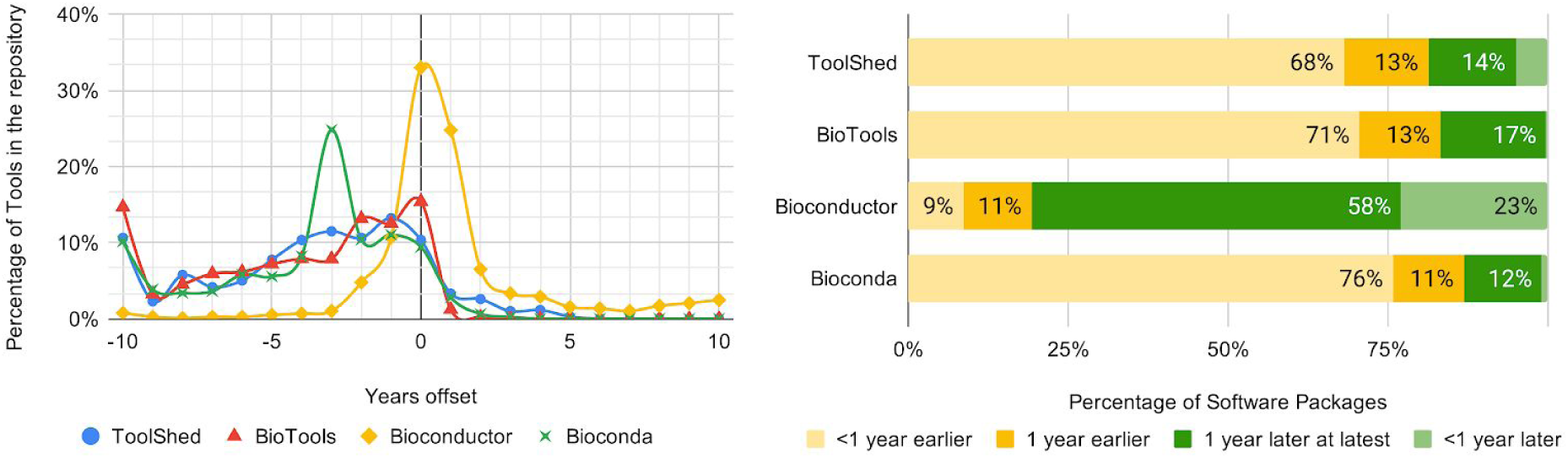
The figure plots the distribution of packages on the year difference between the dates they were uploaded to the PMS, and their respective scholarly paper was published. On the left panel, the negative offset implies that the manuscript was published before its respective software was uploaded to the PMS, and vice-versa for the positive offset. The plots show that 81% of packages are added to Bioconductor before their scholarly paper was published (and 69% within one year apart). In contrast, most packages hosted by Bioconda, BioTools, and ToolShed were uploaded more than one year after their respective scholarly paper was published. Bioconductor’s long establishment, broad community, and maintainer’s commitment to high-quality packages (e.g., they review every package’s source code) can be among the factors that explain such differences in their community norms.

#### Impact on the adoption of their packages

To compare PMS’s impact, we perform a *t*-test on the citation count of publications in relative clusters across the PMS’s. The results of the *t*-test with the null hypothesis that the mean of two populations equals zero are given in Table 3. In general, the tests reject the null hypothesis with a permissive *p*-value or a significant *p*-value with a small Cohen’s *d* effect size for most clusters, suggesting that the PMS’s have a mostly similar impact on the citation count of publications. However, the effect size is large for several mid-cited clusters with a significant *p*-value; for instance, the null hypothesis is rejected with a significant *p*-value and a “very large” effect size comparing the mid-cited publications of Bioconda and BioTools, and BioTools and Toolshed. Therefore, the tests suggest that PMS’s impact on the citation count of high and low-cited publications is mostly similar, while their impact differs for mid-cited publications.

**Table 3.** Results of a two-sided *t*-test for the null hypothesis that the means of citation counts in respective clusters across different PMS’s are identical. With a significant *p*-value, the tests suggest a large difference between the Mid-cited publications of BioTools and Bioconda and BioTools and ToolShed.

| A | B | A's<br>Cluster | B's<br>Cluster | t-Statistic | $p$-value | Cohen's $d$ | Effect size |
| --- | --- | --- | --- | --- | --- | --- | --- |
| Bioconda | Bioconductor | Low | Low | 8.623 | 1.86E-17 | 0.466 | Medium |
|  |  | Mid | Mid | 5.715 | 2.06E-02 | 2.770 | Very large |
|  |  | High | High | 2.373 | 2.78E-02 | 0.531 | Medium |
|  | BioTools | Low | Low | 1.387 | 1.66E-01 | 0.035 | Very small |
|  |  | Mid | Mid | 14.469 | 6.01E-23 | 2.699 | Very large |
|  |  | High | High | - | - | - | - |
|  | ToolShed | Low | Low | 1.544 | 1.23E-01 | 0.102 | Small |
|  |  | Mid | Mid | 1.394 | 1.66E-01 | 0.226 | Small |
|  |  | High | High | 0.826 | 4.14E-01 | 0.246 | Small |
| Bioconductor | BioTools | Low | Low | 9.858 | 8.11E-22 | 0.233 | Small |
|  |  | Mid | Mid | 4.698 | 5.27E-03 | 1.026 | Very large |
|  |  | High | High | - | - | - | - |
|  | ToolShed | Low | Low | 8.559 | 1.83E-16 | 0.656 | Large |
|  |  | Mid | Mid | 6.258 | 1.80E-02 | 2.686 | Very large |
|  |  | High | High | 2.812 | 1.02E-02 | 0.599 | Medium |
| BioTools | ToolShed | Low | Low | 2.830 | 4.98E-03 | 0.094 | Very small |
|  |  | Mid | Mid | 15.258 | 7.17E-24 | 3.008 | Very large |
|  |  | High | High | - | - | - | - |

## Discussion

The bibliography information of every 23,083 software hosted by the PMS’s is extracted and searched on Scopus (see Method section). 46.7% of 23,083 packages either do not have bibliography information or their referenced publication is not indexed by Scopus; 55.8% and 7.7% of packages reference one and at least two publications, respectively (see Figure 2 and Supplementary Figure 1).

The citation per year of every referenced publication was retrieved from Scopus. The total citation count of the publications before and after the respective software was published to the PMS’s is plotted in Supplementary Figure 3. The publications referenced by packages hosted in Bioconductor are least cited compared to those referenced by packages hosted in Bioconda, BioTools, and ToolShed.

### Is citation count a good metric to quantify software adoption?

There is an empirical assumption that *citations quantify the scientific impact*. Hence most quantitative bibliometric evaluations, such as the *h*-index and impact factor, derive from article-level citation count [28]. However, the accuracy of these metrics for evaluating individuals has been criticized, and several concerns have been raised (e.g., challenges of comparing citation counts between fields)[29]. Additionally, evaluating software popularity leveraging publications is limited to findings that have been successful enough to merit publication. However, most tools may not be published whether the authors were not successful in publication or publication was not on their agenda.

The software has been recognized as a legitimate research product by institutions and funding agencies [30]. However, citing a software package has been associated with several open challenges, such as authorship and lack of referencing standards [31,32]. Additionally, the citation of software (e.g., a link to source code) is not tracked by bibliography databases.

The number of times a software was downloaded is a sensible metric to evaluate its popularity. However, the download statistics per year are not available from all the PMS’s studied here. Therefore, the citation count has been used to quantify the adoption of software packages.

### The positive impact of PMS’s on scholarly recognition

We study the citation count pattern of 18,140 publications referenced by 23,083 software packages hosted by software package management systems (PMS) devoted to the Bioinformatics community: Bioconda [22], Bioconductor [23], BioTools, and ToolShed [24]. The study’s objective is to assert if the citation count of publications is boosted after their respective software was published to the PMS’s. We grouped the publications in low-, medium-, and highly-cited clusters and studied the changes in their citation increase pattern separately.

Our results suggest that publications receive a boost citation count after their respective software is published to PMS’s. The impact of the boost is in a similar magnitude to the initial citation count increase. The boost is more considerable for publications with medium citation count than the low and highly-cited articles. Our results suggest that PMS’s impact on the citation count per year rate is in the same magnitude across the studied PMS’s.

The implications of PMS’s on the scientific community goes beyond scholarly paper recognition and software adoption. PMS’s standardized software package installation facilitates their integration with other software packages for developing larger data analysis pipelines. By implementing unified discovery means, the PMS’s facilitate searching for software packages for particular applications. Hence they can reduce redundancy in software package development. Therefore, disseminating software via PMS benefits both individuals and their respective communities.

## Supplementary Material

### Methods

#### Bibliographic databases

There are several bibliographic databases for scholarly articles [6,33]. Among them are Web of Science (WoS), Scopus, and Google Scholar. WoS and Scopus provide subscription-based access to databases of articles published in journals, books, and conference proceedings (see [34–40] for their coverage and accuracy). Google Scholar is primarily a free search engine for scholarly literature that indexes theses, preprints, and technical reports in addition to journals, books, and conference proceedings (see [41,42] for its coverage and quality). The databases are compared in [43–45].

We use bibliographic databases to obtain citation information, associate software across package management systems, and validate free-form user-entered bibliographies of software packages. Google Scholar does not offer programmatic access to its content, and crawling its Web interface violates its End User License Agreement (https://policies.google.com/terms?hl=en). Scopus and WoS both offer programmatic access for subscribed users. Since our institute (Oregon Health and Science University) currently has a Scopus subscription, we used Scopus as the citation information source.

#### Data Acquisition and Cleansing

The data used in the present study is obtained by crawling different package management systems, retrieving software and their scholarly paper’s bibliography information, and retrieving their citation count per year from Scopus. For this purpose, multiple *crawlers* have been developed for the package management systems (see Supplementary Figure 5). The source code is freely available from https://github.com/Genometric/TVQ. In general, the crawlers first obtain a list of the software hosted by the package management systems and then retrieve their details and scholarly information using a specific package management approach. Since the extracted bibliography information was entered mostly in free-text format by users, they do not necessarily adhere to standards. Therefore, we have developed methods that parse the bibliography information and extract their entries when they do not adhere to standards or contain LaTeX-specific typesetting that is not supported by Scopus. For instance:

- @article{id, title={a_title}, doi={NaN}} “NaN” is not a DOI, instead of “NaN” the field should have been removed.
- @article{id, title={a_title}, doi={https://doi.org/10.1109/TKDE.2018.2871031}} DOI should not have a “https://doi.org/” prefix.
- @article{id, title={{a_title}}, doi={{10.1109/TKDE.2018.2871031}}} A single pair of curly brackets should surround each field’s value, but two curly brackets are used here.
- @article{, title={my {<u>\\it</u> title}, author={last_nam{{\\\”e}}}} The title and author fields contain LaTeX typesettings that are not recognized by Scopus.

The method developed to extract bibliography information from free-form BibTeX entries is freely available from the following Github repository as a library: https://github.com/Genometric/BibitemParser

In the following, the crawler of each package management system is explained in detail.

##### ToolShed

We use the ToolShed installation maintained by the Galaxy team, accessible from: https://toolshed.g2.bx.psu.edu

The list of tools hosted by the ToolShed instance is available from the following API endpoint: https://toolshed.g2.bx.psu.edu/api/repositories

This endpoint returns a JSON object containing detailed information for each software package. Among them were their maintainer/owner name, package name, and the date when the package was added to ToolShed. We then use this info to download the package’s zip archive from the following URL: https://toolshed.g2.bx.psu.edu/repos/{owner}/{tool}/archive/tip.zip

The archive contains the package and its Galaxy *tool wrapper*. A wrapper is an XML file that contains a block named “citations”; this XML element contains either the DOI of the package’s respective scholarly paper or their bibliography information presented in Bibtex format. Please refer to the wrapper’s documentation at the following link: https://docs.galaxyproject.org/en/latest/dev/schema.html

##### BioTools

A list of tools hosted by https://bio.tools is available from https://github.com/bio-tools/content. This PMS provides tool metadata in JSON format. The BioTools crawler downloads the repository archive, traverses its “content” folder and parses the JSON file. The JSON file contains the package’s metadata, including its name and its respective scholarly paper’s bibliography information.

##### Bioconductor

We use the current latest list of Bioconductor packages that includes 2,008 tools. To get citation information, we use the Bioconductor software package version 3.10, and first, install the tool and then use the “citation” method to get citation information. We have developed scripts written in Python and R programming languages to extract package information, in addition to their scholarly paper’s citation. The scripts are freely available from the following link: https://github.com/Genometric/TVQ/tree/master/data/bioconductor

##### Search on Scopus

Possible sources to obtain citation information are Google Scholar, Scopus, and Web of Science. Google Scholar is freely available but does not offer public API access; both Scopus and Web of Science are subscription-based. We use Scopus to obtain citation information since our institute has a subscription.

We use Scopus with a limit of 20,000 API requests per week. For this, we developed a wrapper for the Scopus’s “Citation Overview” RESTful metadata API in .NET Core. The API endpoint is: https://api.elsevier.com/content/abstract/citations/

The Scopus search API returns entries that match a given query and may not return any entries if the query slightly differs from the indexed information. For instance, the API does not return any entries for a querying manuscript with the title “A time-varying method for microRNA-mediated sub-pathway enrichment analysis” because the title of the published manuscript contains “subpathway” instead of “sub-pathway.” Alternatively, querying a manuscript with the title “MethylAid: visual and interactive quality control of large Illumina 450k data sets” does not return any results as the published manuscript’s title contains “dataset” instead of “data set.” Alternatively, “Software for the Integration of Multi-omics Experiments in Bioconductor,” because the published title has “multiomics” instead of “multi-omics”. Or: “f-scLVM: Scalable and versatile factor analysis for single-cell RNA-seqR” because of the trailing ‘R’.

When DOI is given, we search using it instead of title; however, DOI is not always given. Additionally, there are some inconsistencies with DOIs as well. For instance, a package references a publication with DOI 1093/bioinformatics/btr345; however, the indexed manuscript’s DOI is 10.1093/bioinformatics/btr345 (note the “10.” in the beginning). Hence Scopus does not return any hits. Such small inconsistencies between the bibliography information in uploaded packages and the indexed literature may cause missing some publications for several software packages.

##### Data Cleansing

The bibliography information of software packages is mostly entered in a free-text format. Bioconductor performs a syntax check on the bibliography information they host; however, not all package management systems perform syntax checks.

Additionally, the values of entries in BibTeX items can be outdated or invalid. For instance, a package owner/maintainer uploads the package with a pre-publication manuscript title. The manuscript may publish with a slightly different title; however, the maintainer may not return to the package management system to update their bibliography entry. For instance, a package is uploaded with the manuscript “A_Stat_Package: Statistical analysis of data”, which was then published as “A_Stat_Package: a tool kit for standardized statistical analysis”.

While some LaTeX typesetting is valid for BibTeX items, they need to be removed before being searched on Scopus. For instance, “Alternative mapping of probes to genes for {<u>\\it</u> Affymetrix} chips” where “{<u>\\it</u> …}” makes “Affymetrix” italic. Scopus does not return correct results in such LaTeX typesettings that are not removed from search queries. The BibItemParser library we developed (https://github.com/Genometric/BibitemParser) removes every LaTeX typesetting when parsing BibTeX items.

Some packages mistakenly cite publications that have been useful to the package’s development. For instance, a tool cites “Least Squares Quantization in PCM”, published in 1982 with ∼11,000 citations, not the publication where the tool is discussed. Proper cleaning would be manually checking every software package’s cited publication, which is impractical due to the number of software packages. Instead, our heuristic excludes publications that are more than 20 years old (i.e., published before 2000 Gregorian calendar).

---

**Supplementary Figure 1.**
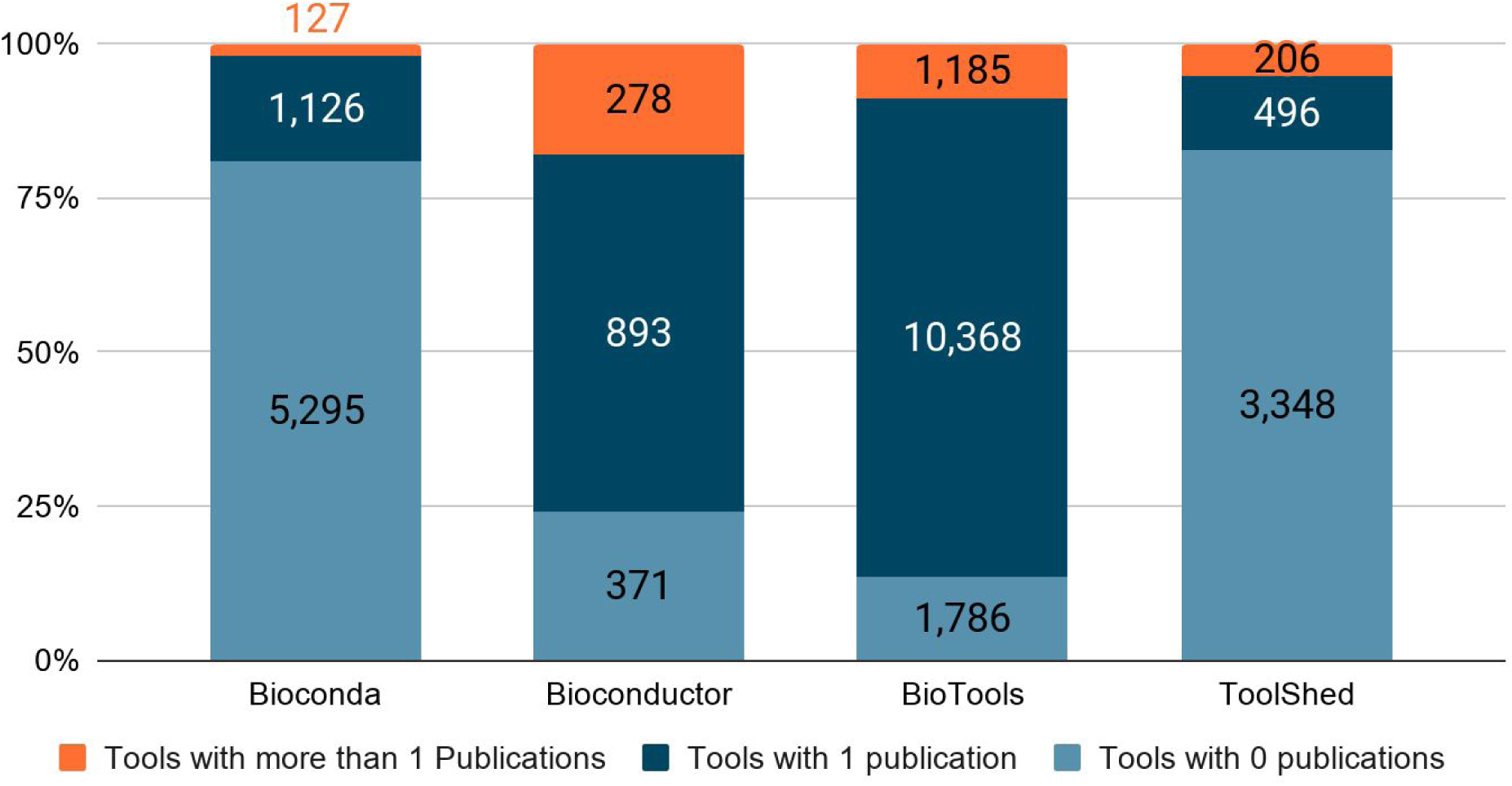
The number of software packages associated with 0, 1, and at least two publications in each package management system. ToolShed and Bioconda contain the most number of packages not associated with any publications (82% and 80% respectively), while most packages in BioTools reference at least one publication (86%).

**Supplementary Figure 2.**
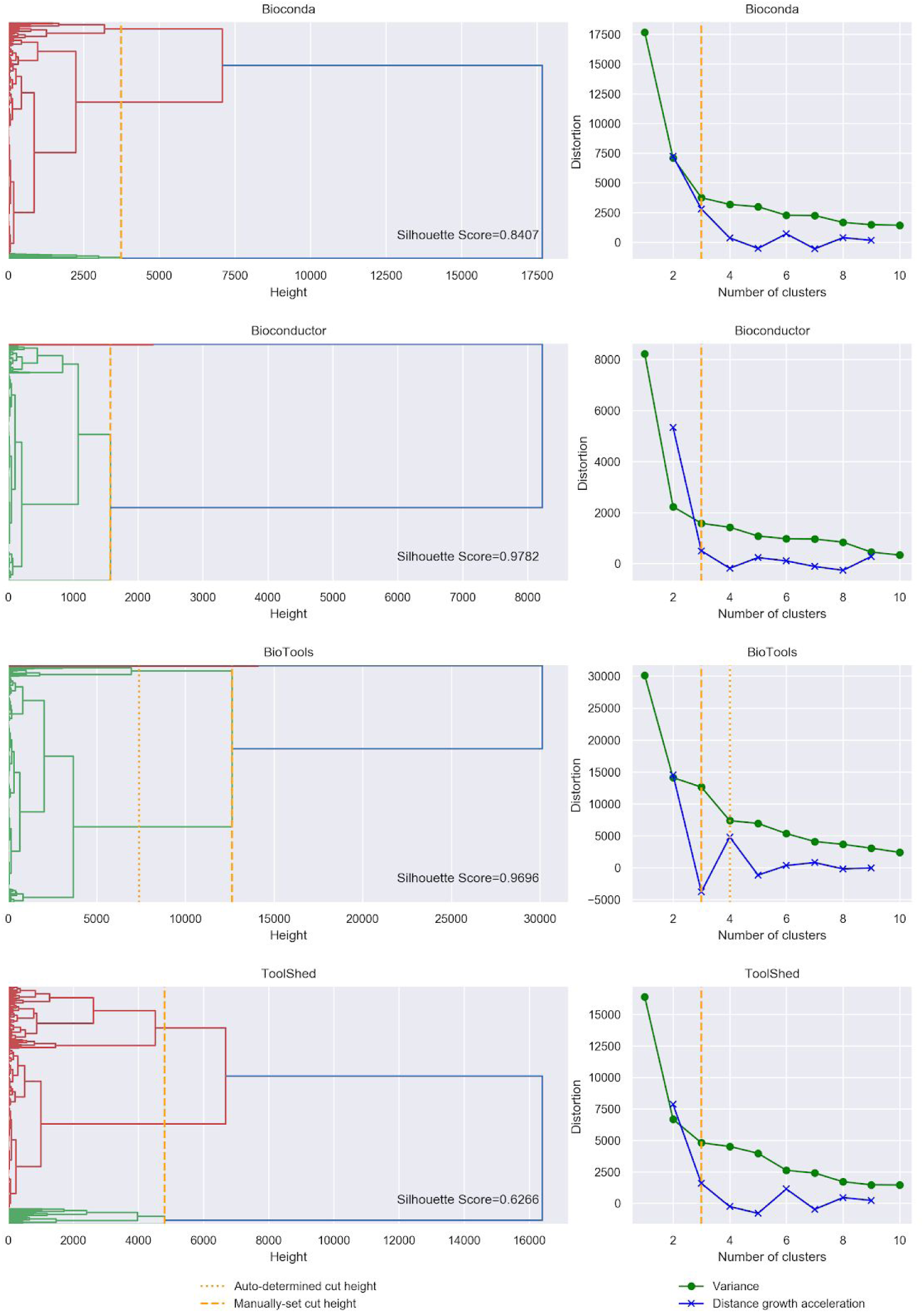
The dendrogram shows the agglomerative hierarchical clustering of tools in different repositories. As is suggested by the dendrogram and the elbow criterion (right panel plots), tools can be grouped in 3-6 clusters. To simplify studying tools in corresponding clusters, we choose to group the tools in 3 clusters, which maintains high cohesion as suggested by the silhouette score of 0.8, 0.9, and 0.9, and 0.8 for Bioconda, Bioconductor, BioTools, and ToolShed, respectively.

**Supplementary Figure 3.**
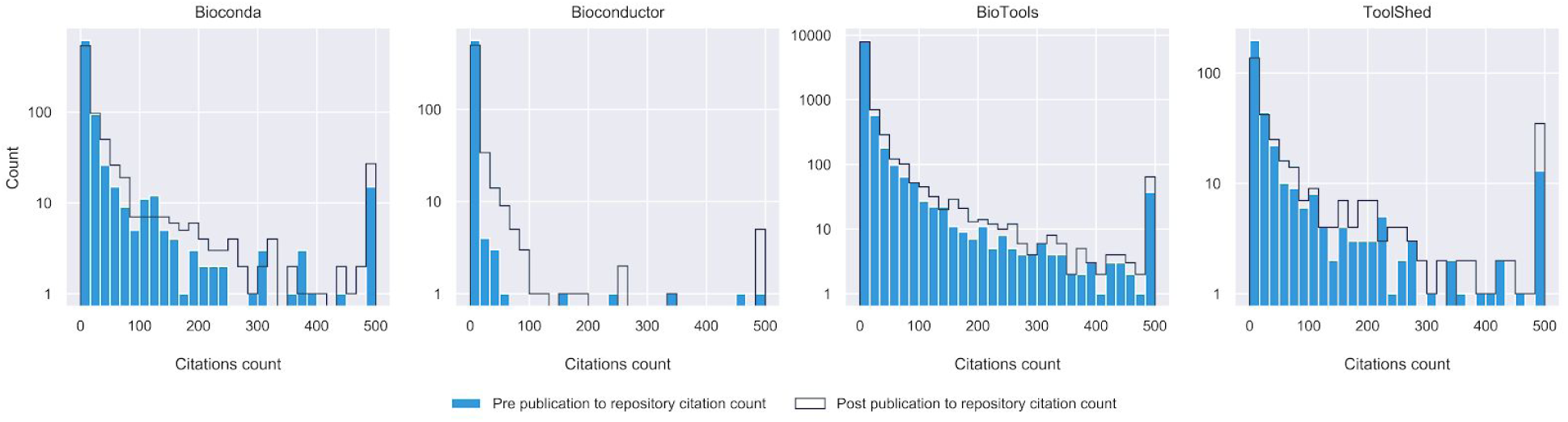
The histogram suggests that the publications referenced by packages hosted in Bioconductor are generally least cited w.r.t those hosted in Bioconda, BioTools, and ToolShed.

**Supplementary Figure 4.**
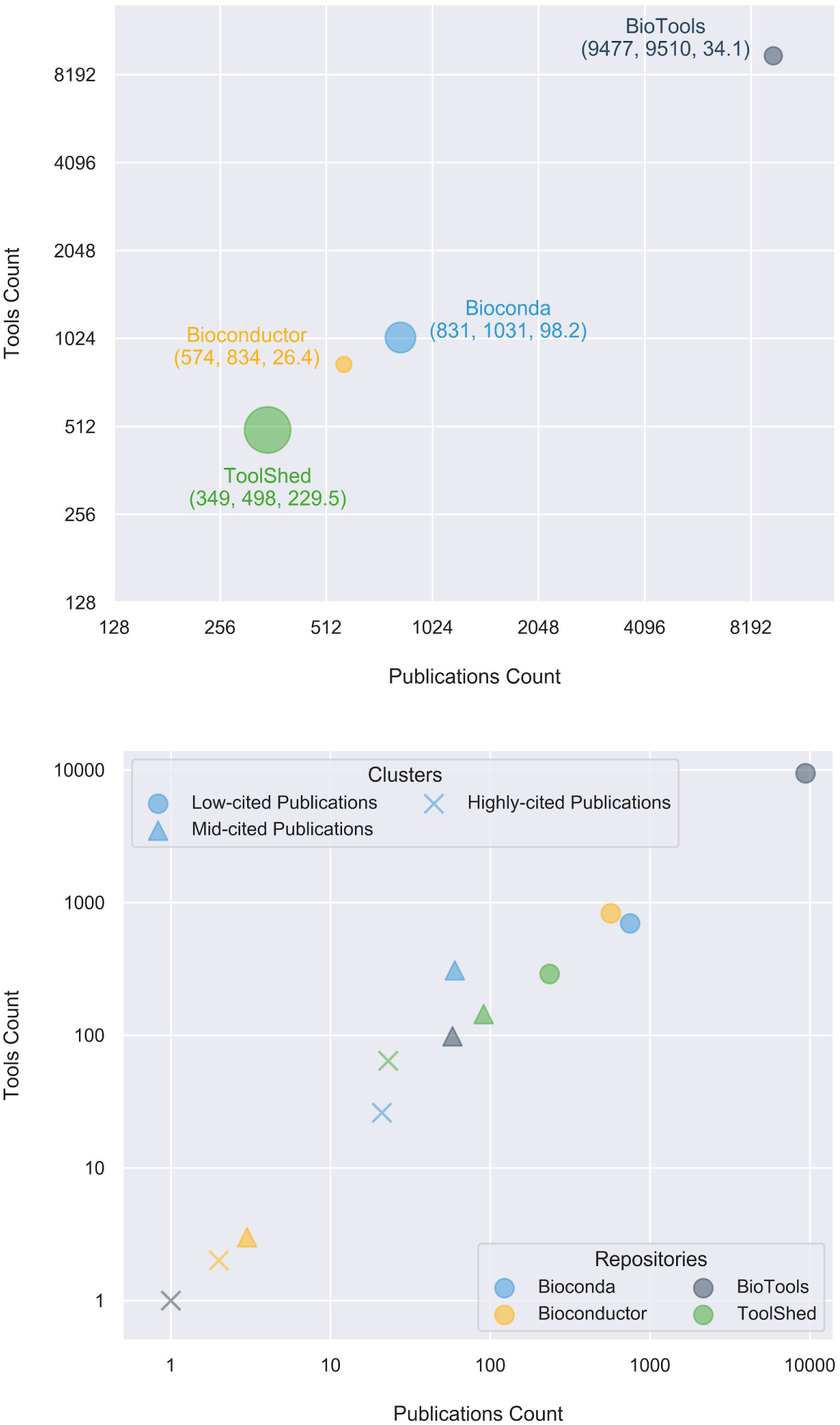
Upper panel: an alternative to Figure 2 where only the tools with at least one publication are included. Bottom panel: the information on the upper panel is divided into clusters.

**Supplementary Figure 5.**
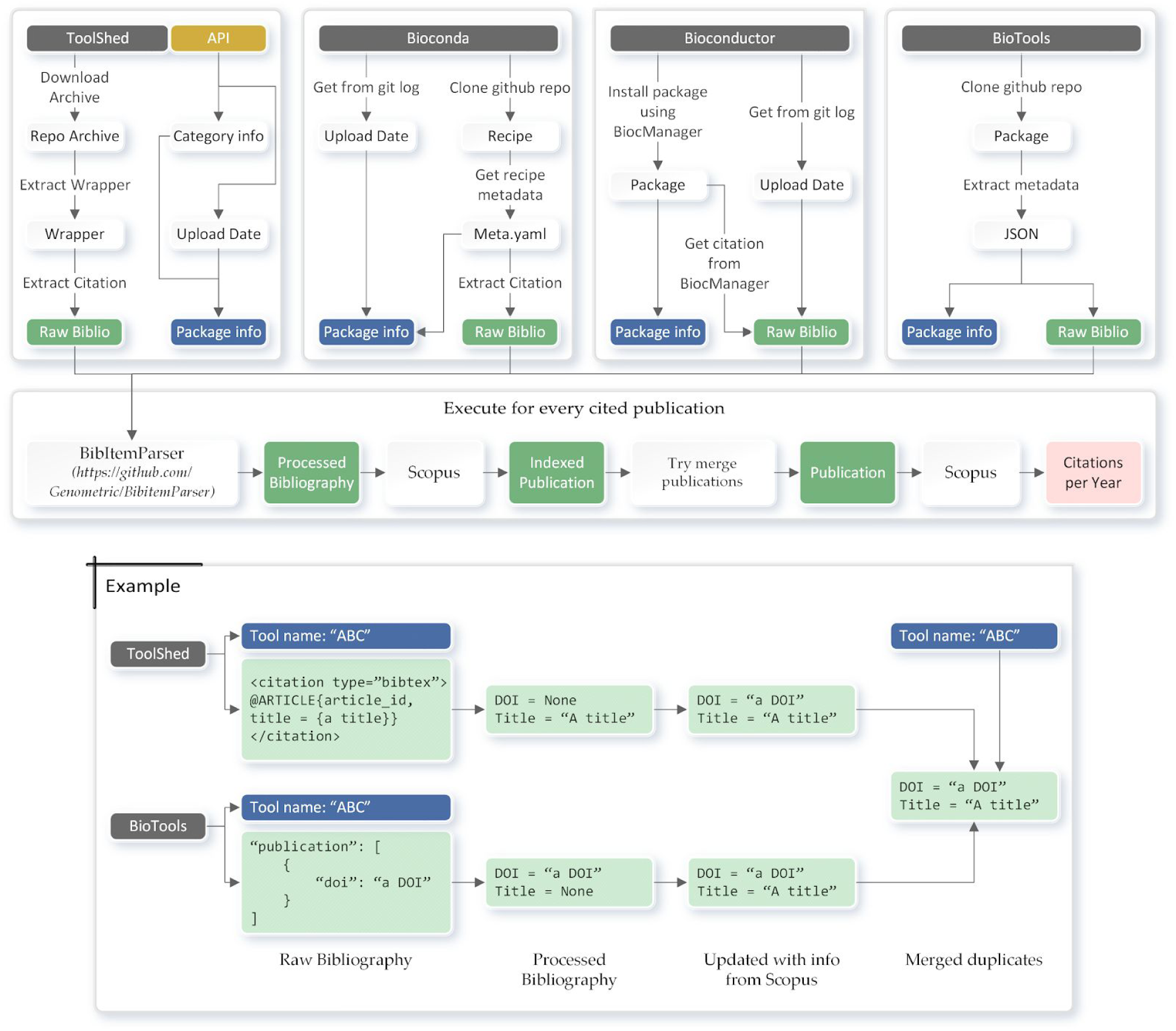
The figure illustrates the process of extracting software and its referenced publication’s information from each of the package management systems and the flow to retrieve citations per year from Scopus. Additionally, it illustrates the process of merging related publications with an example.

**Supplementary Figure 6.**
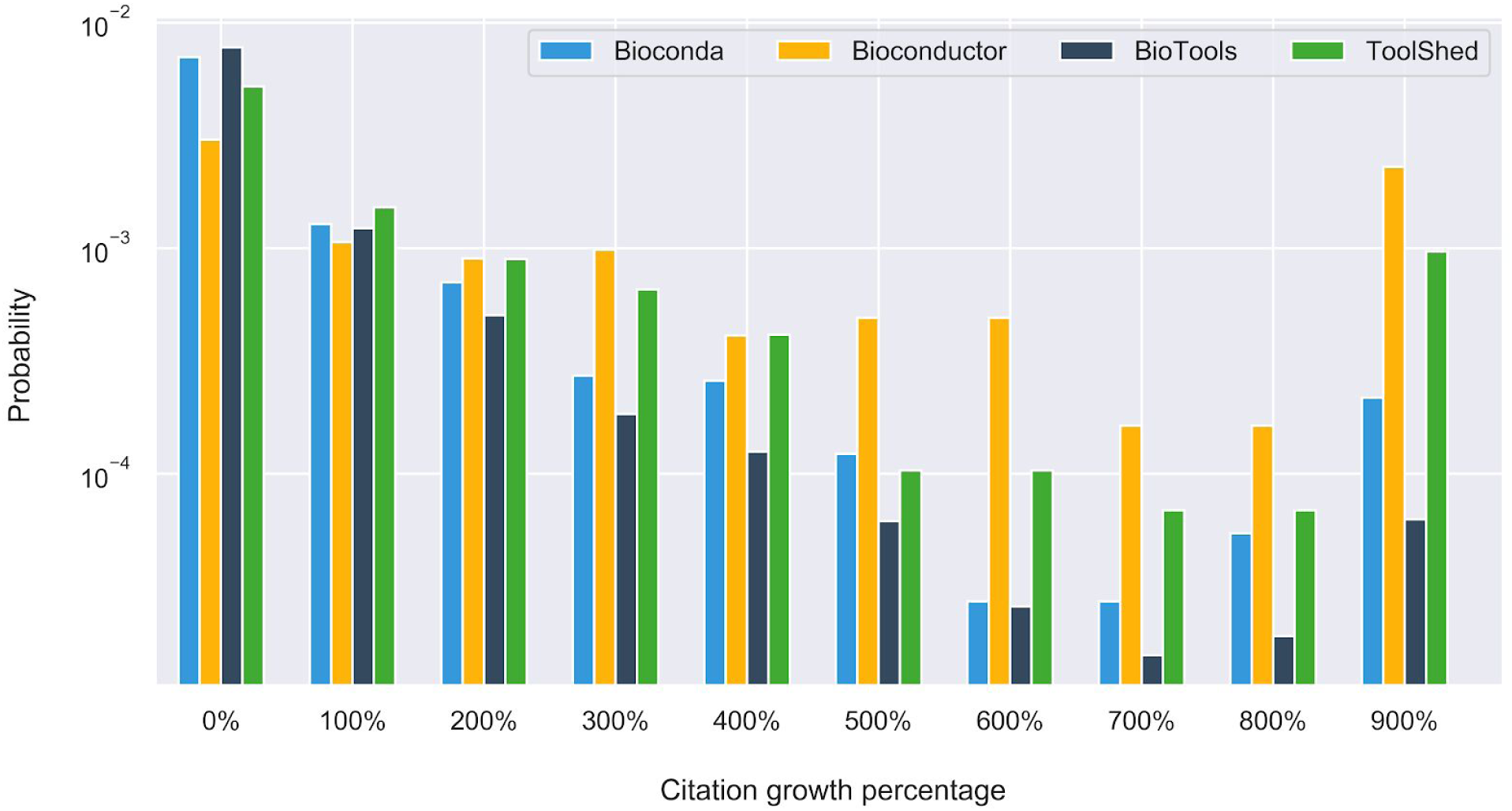
The histogram plots the distribution of the percentage of citation growth, comparing the number of citations before and after a tool was added to the PMS. The rare citation growth cases beyond 1,000% (that span up to 108,000%) are aggregated at 900%. The plot shows that many repositories’ tools receive more than 100% growth in their citation count after adding to the PMS. However, their number diminishes more rapidly for tools in BioTools and Bioconda compared to Bioconductor and ToolShed.

**Supplementary Figure 7.**
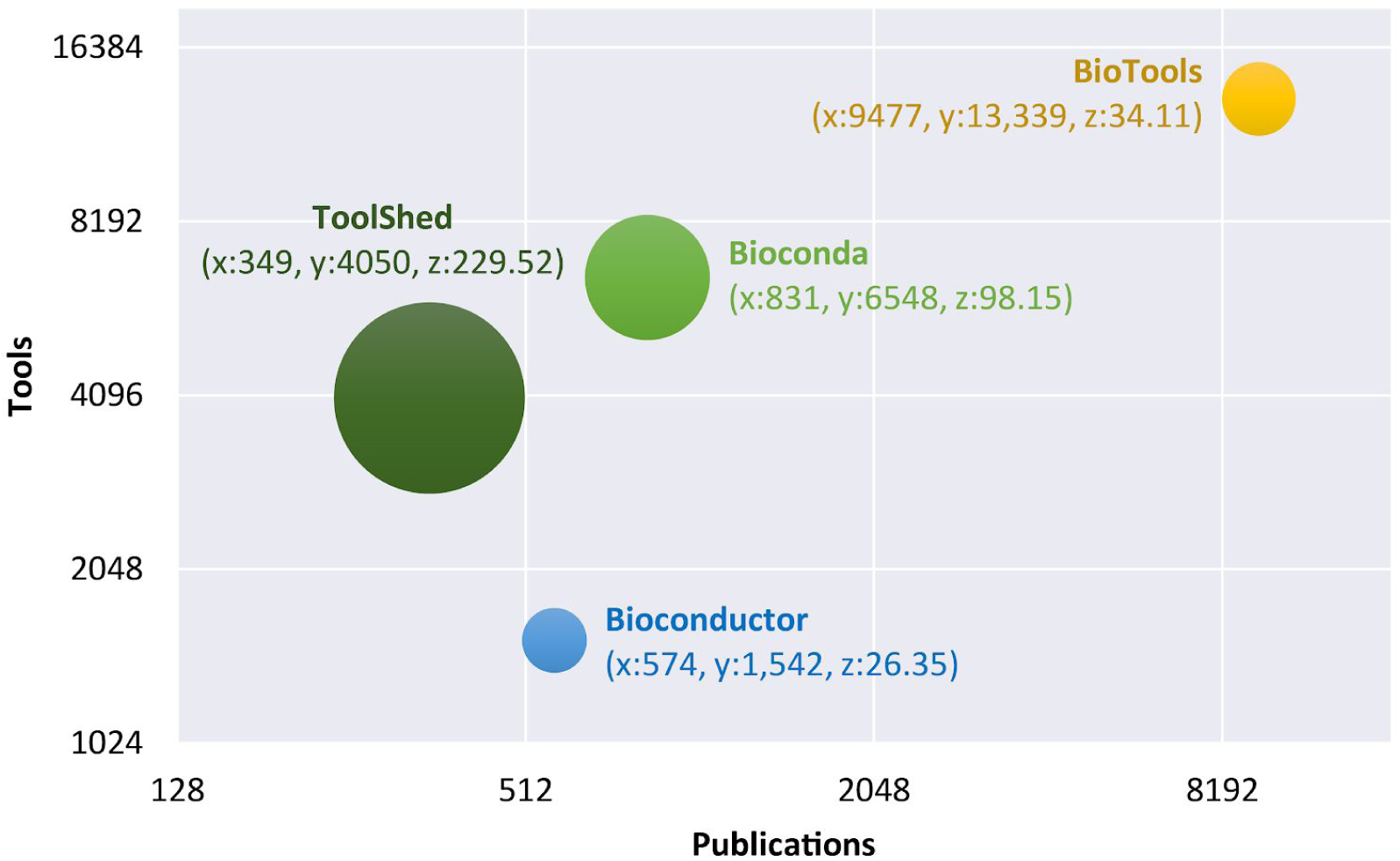
The figure plots the number of packages in a PMS against the number of publications, and the bubble size is proportionate to the average citation count per publication. The plot suggests that while ToolShed has the fewest publications associated with its packages compared to other PMS’s, they are more cited than the publications of other PMS’s publications. In other words, ToolShed contains more popular software w.r.t other PMS’s. Additionally, the plot suggests that the relation between software packages and publications is not 1-to-1, and a significant number of packages reference joint publications; see supplementary figures 1 and 4.

**Supplementary Table 1.** The table contains the average citation count of all publications in each cluster before and after their respective software was published to the package management systems.

|  | Cluster | Average<br>Before | Average<br>After |
| --- | --- | --- | --- |
| Bioconda | Low | 93.3 | 146.9 |
|  | Mid | 1,217.7 | 2,538.3 |
|  | High | 6,837.2 | 12,126.8 |
| Bioconductor | Low | 20.2 | 85.2 |
|  | Mid | 1,667.3 | 10,692.3 |
|  | High | 7,430.5 | 19,516.0 |
| BioTools | Low | 97.5 | 139.9 |
|  | Mid | 6,300.1 | 10,500.8 |
|  | High | 26,819.0 | 31,174.0 |
| ToolShed | Low | 92.2 | 188.2 |
|  | Mid | 1,006.4 | 2,540.1 |
|  | High | 5,754.1 | 11,888.8 |

